# Comparative study of population genomic approaches for mapping colony-level traits

**DOI:** 10.1101/479600

**Authors:** Shani Inbar, Pnina Cohen, Tal Yahav, Eyal Privman

**Author notes:** Equal contribution (Shani Inbar, Pnina Cohen).

## Abstract

Social insect colonies exhibit colony-level phenotypes such as social immunity and task coordination, which are the sum of individual phenotypes. Mapping the genetic basis of such phenotypes requires associating the colony-level phenotype with the genotypes in the colony. In this paper, we examine alternative approaches to DNA extraction, library construction and sequencing for genome wide association (GWAS) studies of colony-level traits. We evaluate the accuracy of allele frequency estimation in sequencing a pool of individuals (pool-seq) from each colony in either whole-genome sequencing or reduced representation genomic sequencing. Based on empirical measurement of the experimental noise in sequencing DNA pools, we show that whole-genome pool-seq is more accurate than reduced representation pool-seq. We evaluate the power of the alternative approaches for detecting quantitative trait loci (QTL) of colony-level traits by using simulations that account for an environmental effect on the phenotype. Our results can inform experimental designs and enable optimizing the power of GWAS depending on budget, availability of samples and research goals. We conclude that for a given budget, sequencing un-normalized pools of individuals from each colony achieves greater QTL detection power.

## Introduction

Social insect colonies depend on the collective performance of a large number of individual workers. Such collective performance gives rise to extended, colony-level phenotypes such as social immune responses to pathogens and parasites, coordinated defense against intruders, information transfer and processing, a colonial pheromonal odor that facilitates nestmate recognition, etc. These colony-level phenotypes are the collective output of individual phenotypes and are affected by both the internal state of the colony, as well as other factors in its environment (e.g., neighboring colonies, predators and competitors). Due to this multilevel complexity, identifying genes responsible for these higher order traits is a challenging endeavor that warrants a multilevel modeling.

In genome-wide association studies (GWAS) genetic variants are associated with phenotypic traits. High-throughput technologies allow genotyping many thousands and even millions of polymorphic markers, most commonly single nucleotide polymorphisms (SNPs). The traits are often quantitative traits, and the genetic elements determining them are known as quantitative trait loci (QTLs). In GWAS, natural populations are sampled and it is generally assumed that individual samples are not related, unlike QTL mapping, in which offspring from controlled crosses are used. Each individual sample is genotyped and phenotyped and a large number of polymorphic loci can be tested for an association with the phenotype of interest. This standard scheme becomes more complicated when one wishes to map colony-level traits because each colony consists of multiple individual genotypes whereas the phenotype is a single measurement at the colony level. Therefore, the methodology should be adapted and take into account this added level of complexity.

Next generation sequencing methods allow for highly cost-effective genotyping. However, sequencing whole genomes in large numbers is still cost prohibitive despite the continuing and dramatic reductions in sequencing costs over the last decade. A project’s sequencing budget is still an important limiting factor because sufficient statistical power for QTL detection often requires many thousands of samples (Meaburn et al., 2005). Consequently, several sequencing-based approaches were developed to reduce per-sample costs. One way to reduce library construction and sequencing costs is pool-seq: pooling DNA from several individual samples and then constructing and sequencing a single library. Multiple pools are constructed, and a locus may be tested for an association of the phenotype of interest with the allele frequency in each pool. In many studies, library construction can amount to over 50% of the total costs. In such cases, pool-seq becomes an especially attractive approach as it allows for an increased accuracy-to-cost ratio for determining a population’s allele frequency (Gautier et al., 2013; Schlötterer, Tobler, Kofler, & Nolte, 2014).

Allele frequency estimation by pool-seq may suffer from experimental noise and biases in various stages of the protocol, including DNA pooling, library construction, sequencing, and data analysis. Van Tassell et al. (2008) assessed the concordance of individually sequenced samples with allele frequencies estimated by sequencing their pooled DNA, and reported a correlation coefficient *rho* = 0.67 (this study used reduced representation genomic sequencing, see below). Futschik and Schlötterer (2010) used simulation studies to compare the power and accuracy of SNP discovery and allele frequency estimation. They simulated allele frequencies of pool-seq and individual genome sequencing over a wide range of parameters and experimental designs and evaluated the effect of unequal representation of individual DNA in pools with population statistics such as Tajima’s π and Watterson’s θ. They concluded that lager pool sizes increase the accuracy of pool-seq, whereas here we report that based on empirical measurements pools of 10 individuals give similar results to pools of 30 individuals. However, these simulations did not take into account potential errors and biases in the representation of individual genotypes in the pool, library construction and sequencing. Such errors and biases are evident from the abovementioned study by Van Tassell et al. (2008), who empirically measured allele frequencies in pool-seq. Moreover, neither of these studies investigated the implications for the probability of detecting QTL when using pool-seq as opposed to individual sequencing in the context of GWAS.

The application of pool-seq to mapping quantitative traits was investigated in multiple theoretical and empirical studies (Bastide et al., 2013; Darvasi & Soller, 1994; Dunnington, Haberfeld, Stallard, Siegel, & Hillel, 1992; Giovannoni, Wing, Ganal, & Tanksley, 1991; Korol, Frenkel, Cohen, Lipkin, & Soller, 2007; Michelmore, Paran, & Kesseli, 1991; Plotsky et al., 1993). These studies used pools of samples taken from the tails of the phenotype distribution in a natural population. They demonstrated that greater statistical power may be obtained with a much reduced sequencing budget when using pool-seq. For example, Bastide et al. (2013) used pool-seq to identify two QTLs for a highly heritable pigmentation trait in *Drosophila melanogaster*. They used simulations to show that their approach is mainly suitable for detecting QTLs with high frequency alleles and reasonably large-effect. When mapping colony-level traits (e.g. of social insect colonies), there is a different motivation to sequence pools of individuals, that is to use of a pool for representing the genotypes of a colony. In these cases, genotypes of individual members of the colony can be assayed as a ‘collective genome’, which is then tested for association with the collective phenotype.

Another way to reduce the per-sample costs is reduced representation genomic sequencing approaches, such as Restriction-site Associated DNA sequencing (RAD-seq). In RAD-seq (also known as genotyping-by-sequencing; GBS), restriction enzymes are used to limit the sequencing effort to a subset of randomly distributed loci in the genome (Baird et al., 2008). This approach has several advantages. First, it reduces the amount of sequencing needed per sample by an order of magnitude or more. Second, it can be used to carry out population genetic studies without a reference genome, making it applicable to non-model organisms with very low sequencing budgets. Third, unlike SNP microarrays that need to be designed before their use, RAD-seq discovers and genotypes SNPs simultaneously in a single sequencing run. Fourth, the costs of homemade RAD-seq library construction can be 5-10 fold lower than those of commercial whole-genome shotgun libraries (Peterson, Weber, Kay, Fisher, & Hoekstra, 2012). RAD-seq, therefore, enables construction and sequencing of individually indexed libraries for thousands of samples at as low as 10 USD per sample.

Fifth, by using enzymes with different cutting frequencies (4-, 6- or 8-cutters) RAD-seq allows trading off a smaller number of loci sequenced for a larger number of samples or greater sequencing depth (depth is important for accurate genotyping). Thus, RAD-seq protocols can be tailored to particular genomes and different studies by adjusting the sequencing depth and the proportion of the genome that is sequenced according to scientific goals. For example, when choosing a sequencing approach for a genomic mapping study, one should take into consideration the rate of decay of linkage disequilibrium (LD). RAD-seq would be beneficial for QTL mapping using closely related samples as markers would be linked across many kilobases. However, when using unrelated individuals for GWAS, LD decays much faster, and may not be detectable even for markers less than 1 Kbp apart (depending on the rate of recombination in the study organism). In these cases whole genome sequencing may be necessary. It has also been argued that RAD-seq is not suitable for other types of genomic scans, such as detection of loci involved in local adaptation (for an interesting discussion see (Andrews, Good, Miller, Luikart, & Hohenlohe, 2016; J. M. Catchen et al., 2017; Lowry et al., 2017a, 2017b; McKinney, Larson, Seeb, & Seeb, 2017).

Here we describe a methodological study evaluating the contribution of alternative approaches to the statistical power of GWAS for mapping colony-level traits. We applied whole-genome sequencing, RAD-seq, pool-seq, and their combination to ant colony samples, providing a broad overview of the expected results of different sequencing approaches. We report the minimum number of individually sequenced samples per colony that is equivalent to a pooled sample for the purposes of GWAS. We also take into consideration environmental factors that affect the phenotype to various degrees (i.e. different levels of heritability) which can substantially reduce the power of the analysis. These results enable optimization of experimental designs for GWAS of colony-level traits and show that pool-seq is a significantly more powerful approach for mapping colony-level traits, although depending on availability of samples and scientific goals, individual sequencing may be advantageous.

## Methods

### Model organism

Our study population was sampled from Betzet beach on the northern Israeli coastline. This is a population that was previously described as *C. drusus* (Eyer, Seltzer, Reiner-Brodetzki, & Hefetz, 2017), but our recent species delimitation study raised the question of whether *C. drusus* is separate species or is it the same species as *C. niger*, because these populations are not differentiated by their nuclear genomic DNA (Reiner-Brodetzki, Inbar, Cohen, Hefetz, & Privman). Colonies of this population are monogyne (headed by a single queen), polyandrous (queens are multiply mated) and monodomous (single nest per colony) (Eyer et al., 2017). We chose a monogyne population in order to avoid substructures of the multiple families within a polygyne colony, which would further complicate the analysis. A monogyne, polyandrous population in this species has a mean within-colony genetic relatedness of 0.26 (Leniaud, Heftez, Grumiau, & Aron, 2011).

### Samples and pooling schemes

Thirty workers were sampled from each of three colonies (colony BZT48: N33.05277, E35.10277 colony BZT49: N33.05243, E35.10278 colony BZT50: N33.05162, E35.10245) and every ant was individually frozen in −80°C on the evening of the same day. Workers of all sizes were included in the sampling. DNA was extracted individually from whole bodies, using TRIzol® (Invitrogen Life Technologies) and kept in −20°C. Quantity and quality of extracts were determined by NanoDrop™ 2000 Spectrophotometer and gel electrophoresis was used to verify sample consistency, i.e., DNA yield, degradation, and phenol/salts residues. Pooled samples were prepared by normalizing DNA quantities to ensure equal representation of every individual sample in the colony’s pooled sample.

### RAD library construction

A reduced representation genomic library was constructed according to a modified double-digest Restriction-site Associated DNA sequencing (ddRAD-seq) protocol, based on protocols from Parchman et al. (2012) and Peterson et al. (2012). Briefly, DNA was digested by two different restriction enzymes (EcoR1 as a rare-cutter; Mse1 as a frequent-cutter) and ligated to barcoded adaptors for multiplexing (using CutSmart buffer® NEB). Amplified products (using Q5 Hot Start Polymerase® NEB, the number if cycles was reduced and the number of replicates and starting DNA volume were increased. Also, primers and dNTPs were added to the final thermal cycle in order to reduce production of single-stranded or heteroduplex PCR products) were separated by gel electrophoresis and insert sizes of 300- 400bp were selected and purified (Qiagen, MinElute Gel Extraction Kit, followed by Beckman Coulter, Agencourt® AMPure® XP). The library was sequenced in one lane of a single-end, 100bp reads, on a HiSeq4000 Illumina sequencer. Sequenced data was submitted to NCBI SRA database (Accession: PRJNA494296).

### Whole genome library construction

In addition, for 29 individuals of colony BZT50, DNA was extracted from three body segments, i.e., heads, thoraces and abdomens, but without legs. DNA was also extracted from all legs pooled together (174 legs). One pooled sample of the 29 individuals was normalized for equal quantitative representation of each sample and two additional normalized pools were independently prepared from only 10 individuals out of the same 29. Uniquely barcoded genomic libraries were constructed for each of these 33 individual/pool samples (Illumina® TruSeq® Nano DNA Library Prep kit). Briefly, with this method, a Covaris S220 sonicator was used for DNA shearing of 550bp inserts, ends were repaired, and the fragments were size-selected and purified using magnetic beads (supplied in kit). Then, an adenosine was added to the 3’ end of the blunt fragments, individually barcoded adaptors were ligated, and ligated fragments were amplified. Libraries were normalized and pooled so that the four pools were represented five-fold more than individual samples. The library was sequenced in one lane of a paired-end, 150bp reads, on a HiSeq4000. Sequenced data was submitted to NCBI SRA database (Accession: PRJNA494296).

### RAD-seq data analysis

Samples were analyzed by STACKS (J. Catchen, Hohenlohe, Bassham, Amores, & Cresko, 2013; J. M. Catchen, Amores, Hohenlohe, Cresko, & Postlethwait, 2011) and Bowtie2 (Alexander, Novembre, & Lange, 2009)to create a catalogue of single nucleotide polymorphism (SNPs). Sequencing was in high quality (Mean phred scores ranged between 30-40 for all bases i.e., a chance of 1 to 1000 for sequencing error at worst case scenarios). The number of raw reads totaled in 302,654,900, of which 178,631,035 were associated with a sample (barcoded). 166,681,127 reads remained after filtering low-quality reads using process_radtags from the STACKS package. The average depth per sample of the retained reads ranges between 0 and 132X, averaging at X13.4 per sample (calculated based on 67,687 restriction sites in the reference genome). Reads were aligned to the *C. niger* reference genome version Cnig_gnA (NCBI accession number pending) and those with low alignments scores or multiple mapping were filtered out, leaving 128,571,843 reads (See Table 1). Remaining reads were analyzed together to create a catalogue of all alleles found using the STACKS pipeline, resulting in a catalogue of 25,590 SNPs.

**Table 1:** Sequencing reads per colony before and after initial processing and after alignment based filtering.

| Colony | Total reads | Retained reads after initial processing | Average retained reads per sample |
| --- | --- | --- | --- |
| 48 | 128,572,232 | 123,053,180 | 3,515,805 |
| 49 | 54,838,668 | 50,950,729 | 1,592,210 |
| 50 | 4,7106,250 | 43,398,504 | 1,496,500 |

### Whole genome data analysis

Sequences were analyzed using Bowtie2 and GATK (McKenna et al., 2010)to create a catalogue of SNPs. Sequencing was in high quality (Mean phred scores ranged between 20-30 for all bases). Number of raw reads totaled in 675,094,588 (2 X 337,547,294) of which 628,874,982 (2 X 314,437,491) were associated with a sample. Duplicated reads were removed and low quality reads were filtered leaving 614,306,762 reads (97%). Low quality reads were trimmed or removed using Trimmomatic software, leaving sequences that were at least 75bp long. Total number of remaining reads was 529,101,399 (average coverage 6.4±2.2X per sample based on genome assembly size of 296,608,662bp). Reads were aligned to the reference genome and 454,441,029 reads remained after filtering low-scoring alignments and reads with multiple mapping (70% of raw reads).

Genotypes of individual samples were inferred by GATK, using the standard approach for diploid genotypes. Briefly, SNP calling was performed using HaplotypeCaller and GenotypeGVCFs from the GATK package, using the Bayesian approach. For pooled samples, the proportions of reads supporting each allele were calculated at each locus. Allele frequencies of the pool sample were compared to the frequencies in the individual genotypes. GATK produced a catalogue of 1,000,801 SNPs and, after removal of indels and SNPs with more than 2 alleles, 864,743 SNPs were retained. SNPs with excessive coverage (>450X total coverage) were removed (suspected repetitive sequences), leaving 863,106 SNPs.

### SNP filtering

We tested a range of parameters configurations for SNP filtering. A minimal number of reads covering a SNP (*m*) is required to call the genotype of a particular sample in a particular SNP locus. We set a higher minimum for the pooled sample (10, 15, 25, or 30) then for individual sample (2, 4, 6, or 8). SNPs were required to have called genotypes for a minimal percentage of individual samples (*r*), ranging between 75 and 95, and a minor allele frequency (MAF) of at least 1.25%, 3%, or 9% (*min_maf*). A total of 400 different filtering options were tested. Spearman ranked correlations were calculated between the individuals and each of the pooled samples. The following filtering criteria were chosen based on the higher correlation obtained between the pooled and the individual genotypes. For the RAD-seq data, filtering criteria were: *r* = 90%, *min_maf* = 3%, *m* = 4 for an individual sample and *m* = 10 for a pooled sample. For the whole genome sequence, filtering criteria were: *r* = 75%, *min_maf* = 1.25%, *m* = 2 for an individual sample and *m* = 10 for a pooled sample.

### Simulations

The purpose of the simulations was to compare the power of GWAS using the pool-seq approach to that of sequencing of individual samples. For a realistic allele distribution we used population allele frequencies of 10,000 SNPs, sampled from an empirical distribution of SNP allele frequencies of 310 RAD sequenced *Solenopsis* fire ants (Privman et al., 2018) Based on the sampled allele frequencies, genotypes for the queens and her mates (five mates per queen) were sampled randomly for 50 monogyne, polyandrous colonies (Figure 1). Then, 30 worker daughters were simulated for each colony based on the parents’ genotypes. Three replicates of pool-seq results (for these 30 workers) were simulated according to the empirically observed relationship between the individual genotypes and the pool allele frequency in the whole genome sequencing data (above). We assumed a single SNP as a QTL (a causal SNP affecting a single quantitative trait) and association tests were done to evaluate power for detecting that QTL. We examined 30 alternative scenarios for mapping the QTL using the genotypes of between 1 and 30 individuals per colony. Additionally, we examined the power of using the average of triplicates of pools from each colony.

**Figure 1:**
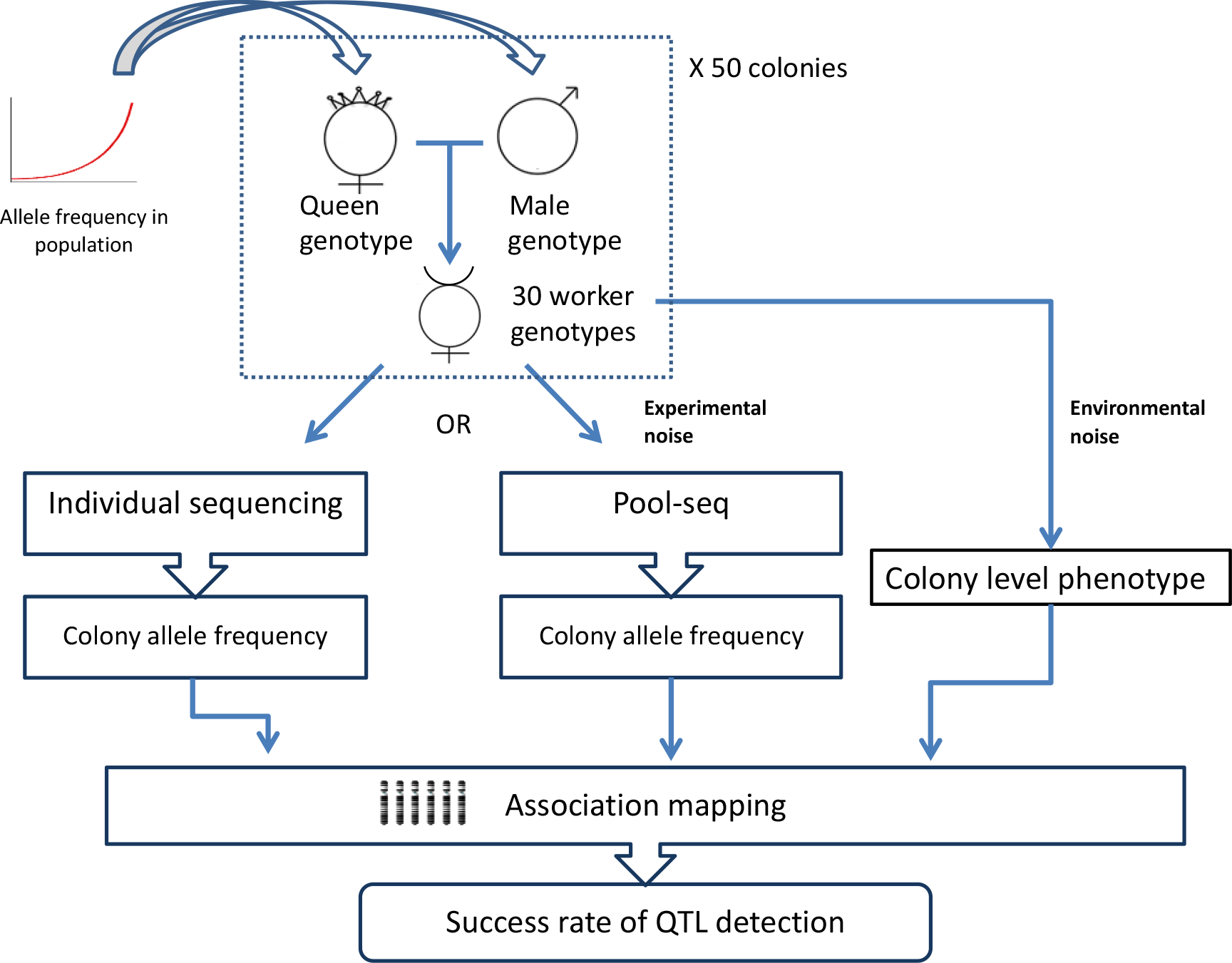
Simulation scheme. Allele frequencies of 10,000 SNPs are sampled from an empirical distribution. Genotypes were randomized for the mother and five fathers of each colony, and then for 30 of their daughter workers. The corresponding pool allele frequencies were sampled according to the empirical correlation between pools and individuals. Colony level phenotypes were generated based on the individual genotypes at a single phenotype-affecting SNP and at varying environmental noise levels. The success rate for detecting the causal SNP was determined for 100 simulated datasets.

Simulation were done under the assumption that there is no linkage disequilibrium between the simulated SNPs. Environmental factors affecting the quantitative trait were accounted for by adding varying levels of “noise” to the phenotype - a randomly chosen number from a normal distribution with a mean of 0 and standard deviation of 0, 0.5, 1, 1.5, 2, 2.5 or 3.

50 colonies were simulated 100 times. Each time, a single SNP was chosen out of the 10,000 SNPs, and 10 phenotypes, for each level of noise, were simulated. In total, 1000 simulated data sets were generated. For each data set, we attempted to predict which of the 10,000 SNPs affects the phenotype in each of the scenarios (1-30 individuals, or a pool), using linear regression. A prediction was determined as successful when the QTL was identified with p-value < 0.05, after correcting the results for 10,000 tests by FDR (Benjamini & Hochberg, 1995).

## Results

### Allele frequencies and their correlations between pool-seq and sequencing of individual samples

Individual samples and three types of pools were analyzed by whole-genome sequencing (WG-seq; see Methods): a pool of 29 individual samples, two pools of 10 samples (out of the same 29), and a non-normalized pool of legs (from the same 29 samples). Depending on filtering criteria, *rho* values ranged between 0.85 and 0.96 (Spearman), but the highest correlation was achieved with very strict filtering criteria that left very few SNPs. Therefore, we chose a more relaxed filtering criteria: minimum number of 2 reads covering a SNP in individual samples and 10 reads in the pool sample, no more than 25% of missing data per SNP, and a minimum minor allele frequency of 1.25%. The different pooled samples showed similar *rho* values; number of SNPs were 643,841-654,398 and *rho* correlation values were 0.899-0.894 (Table 2).

**Table 2:** Correlations between allele frequencies in individual genotypes and pool-seq.

| Colony/pool type | Individual coverage (average and std. dev.) | Pool coverage | Number of SNPs | Correlation |
| --- | --- | --- | --- | --- |
| RAD-seq Colony 48 | 26 ± 38X | 3.7X | 1,656 | 0.681 |
| RAD-seq Colony 49 | 12 ± 10X | 6.0X | 2,161 | 0.749 |
| RAD-seq Colony 50 | 11 ± 11X | 19X | 2,431 | 0.715 |
| Whole-genome with pool-10-1 | 6.4 ± 2.2X | 34X | 653,322 | 0.899 |
| Whole-genome with pool-10-2 |  | 38X | 654,398 | 0.894 |
| Whole-genome with pool-30 |  | 27X | 643,841 | 0.890 |
| Whole-genome with pool-legs |  | 26X | 646,872 | 0.892 |

Similarly, we chose these filtering criteria for the RAD-seq data: a minimum number of 4 reads for individual samples and 10 reads for pooled samples, 10% missing data per SNP, and 3% minimum minor allele frequency. The estimated correlation between pools and individual sequencing was much lower than that of WG-seq. The colony that received the greatest sequencing depth had only a *rho* of 0.749 (colony 49) whereas the lowest *rho* value was 0.681 (colony 48).

For both RAD and WG libraries, a comparison of the allele frequency distributions between individual sequencing and pool-seq data shows that the pool-seq analysis assigns 100% allele frequency to many alleles that have less than 100% frequency in the individual genotypes (Figure 2). Furthermore, there is a gap with very few SNPs between 80% and 100% allele frequency in the RAD pool-seq allele distribution, and between 90% and 100% in the WG pool-seq. Conversely, the individual genotypes show that many SNPs should have allele frequencies in this range.

**Figure 2:**
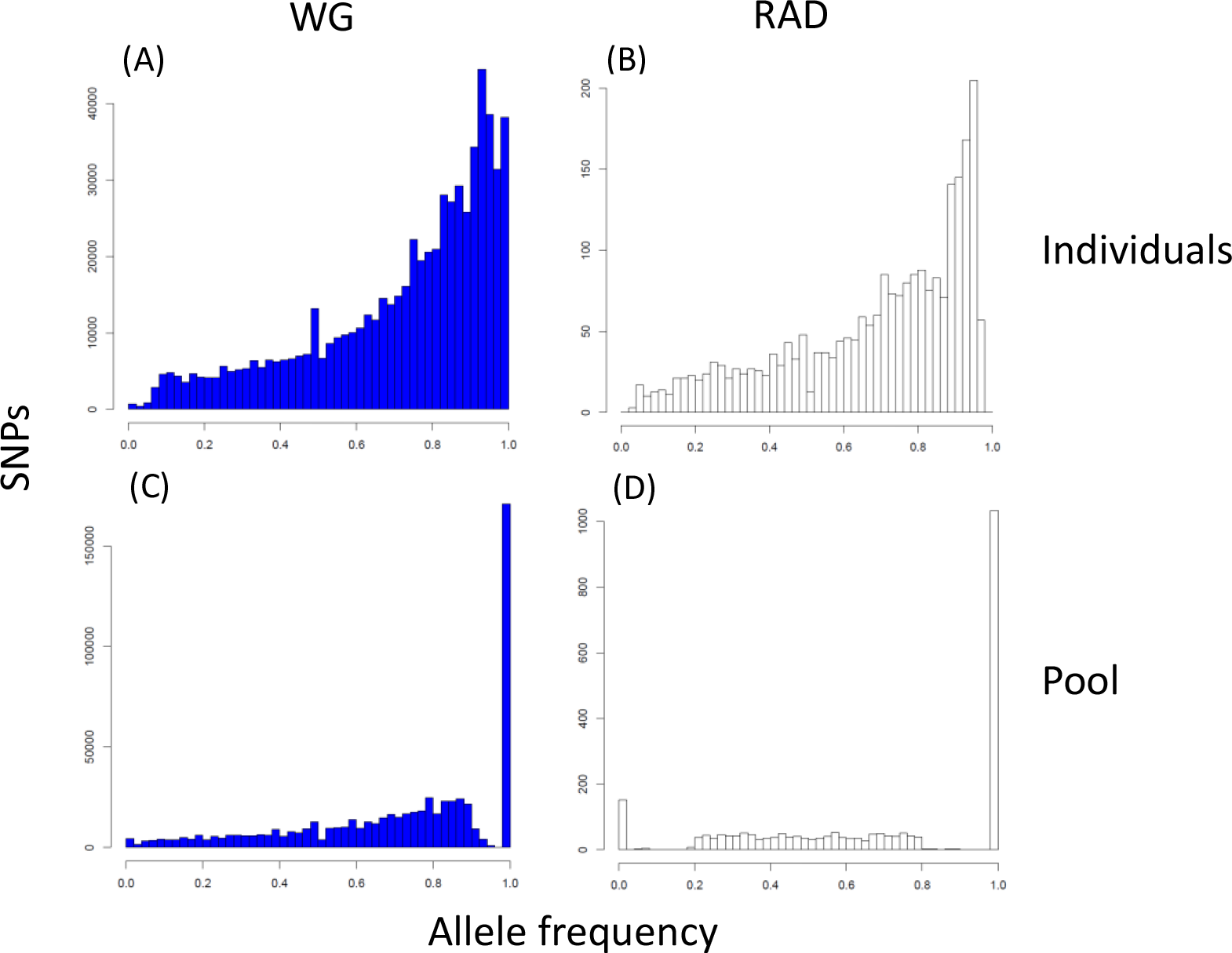
Allele frequency distributions according to whole genome (WG) sequencing and RAD-seq of pooled samples vs. individual samples. **(A&B)** Frequency distribution for the reference allele of 643,841 SNPs in 29 whole-genome sequenced individuals (A) or 30 RAD-sequenced individuals (B). **(C&D)** Allele frequencies in whole genome sequencing (C) or RAD-seq (D) of the pooled DNA from the same 29 or 30 individuals.

### QTL detection power for GWAS using pool-seq

Next, the statistical power of the various approaches for detecting a QTL was evaluated in a simulation study, in which the genotypes of workers from 50 colonies were simulated. One SNP was assumed to be a QTL of a single colony-level phenotype. Various levels of noise were added to the phenotype, to represent an environmental effect on top of the genetic effect of the QTL. A regression test was used to detect the association of the phenotype with the genotypes of a set of individual samples or the allele frequency estimated by pool-seq. When using 30 individual samples, the detection rate decreases from 100% with no environmental noise to 7.1% with noise level of 3, as expected (Figure 3). The noise level corresponds to the size of the environmental effect relative to the genetic effect. For example, a noise level of 1 means that the environmental effect is on the same order of magnitude as the genetic effect of the QTL. Also as expected, success rate increases with the number of individuals used, but it plateaus for five or more individuals at low noise levels and for 12 or more individuals at the highest noise level. With noise level of 1, the power of pool-seq (58.2%) is on par with the power of sequencing five individuals (58.4%). In this case, sequencing six or more individuals per colony would provide greater power than pool-seq. As noise increases, the pool’s power is equivalent to the power of a larger number of individuals. For example, for a noise level of 1 the pool is equivalent to five individuals, while for a noise level of 2 it is equivalent to nine individuals. However, if pools of 150 colonies are used, the power increases dramatically to 92.3% for a noise level of 1 (such numbers would be possible within the same budget of sequencing six individuals per colony for 50 colonies; see cost analysis below).

**Figure 3:**
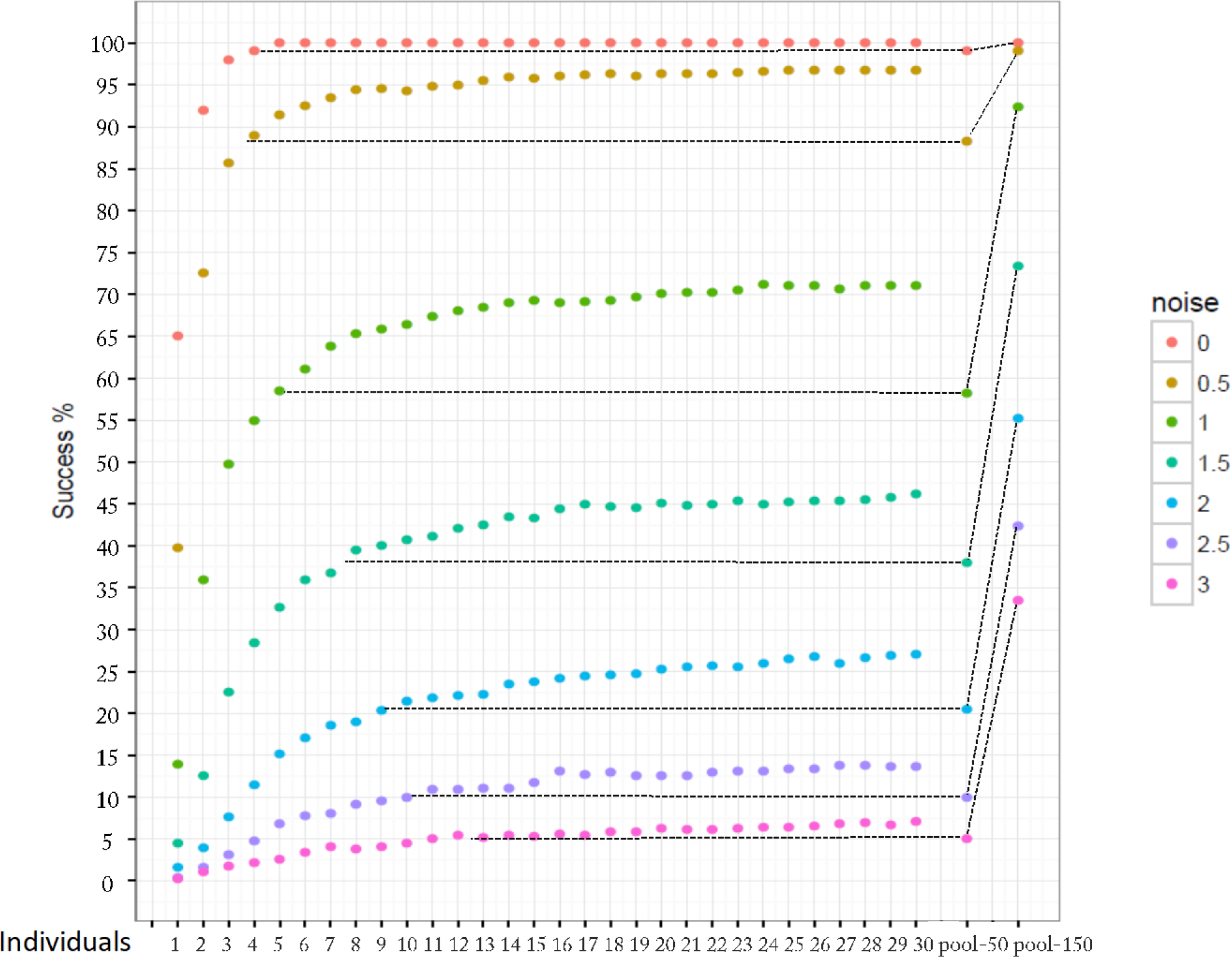
Power of QTL identification. Success rate for detecting a QTL with p<0.05 in 1,000 simulations of 50 colonies (or 150 colonies in the case of the pool-150 results), with individual sequencing of 1 to 30 samples or pool-seq, and with varying levels of environmental noise. Different colors show different levels of noise that affect the phenotype. The dotted lines illustrate the number of individual samples that provides equivalent power as a pool when only 50 colonies are used, whereas 150 colonies provide much greater power as shown by the diagonal dotted lines.

### Cost effectiveness

To aid future studies in determining what would be the favorable DNA extraction, library construction and sequencing approach, we estimated the costs for the various alternatives (Table 3). Sequencing six individuals from each of 50 colonies (by WG-seq) would require a budget of approximately 19500 USD. For the same budget, the number of colonies that can be analyzed using the alternative approaches ranges up to 156 or 1000 colonies for WG-seq or RAD-seq, respectively (when individuals are pooled before DNA extraction). These estimations assumed a desired coverage of 10X per individual sample and 30X for pool sample. The costs include the number of DNA samples extracted and the number of libraries constructed in each approach. The proportion of budget used for each expense component is shown in Figure 4. One may also consider the difference between the alternatives in terms of labor. Extracting DNA and constructing libraries in large numbers is rather labor intensive, and this is not represented here as a cost. This analysis can help future studies determine a combination of sampling, pooling and sequencing approaches which best fit scientific goals.

**Table 3:** The number of colonies which can be included is a GWAS for a given budget, in the alternative approaches.

|  |  |  | Colonies | Samples | Extraction cost | Libraries | Library prep cost | Sequencing cost | Total cost |
| --- | --- | --- | --- | --- | --- | --- | --- | --- | --- |
| **WG-seq** | Individual seq |  | 50 | 300 | 1500 | 300 | 9000 | 9000 | 19500 |
|  | Normalized pool-seq |  | 115 | 1147 | 5735 | 115 | 3441 | 10324 | 19500 |
|  | Un-normalized pool-seq |  | 156 | 156 | 780 | 156 | 4680 | 14040 | 19500 |
| **RAD-seq** | Individual seq |  | 197 | 1182 | 5909 | 1182 | 11818 | 1773 | 19500 |
|  | Normalized pool-seq |  | 302 | 3023 | 15116 | 302 | 3023 | 1360 | 19500 |
|  | Un-normalized pool-seq |  | 1000 | 1000 | 5000 | 1000 | 10000 | 4500 | 19500 |

Cost estimates in USD. Assuming 5 USD per DNA extraction; 30 USD per WG library construction and 10 per RAD library; 10 USD per Gb sequencing; 3 Gb for a genome coverage of 10X per individual sample or 9 Gb for 30X per pool; and 20-fold less sequencing for RAD-seq relative to WG-seq.

**Figure 4:**
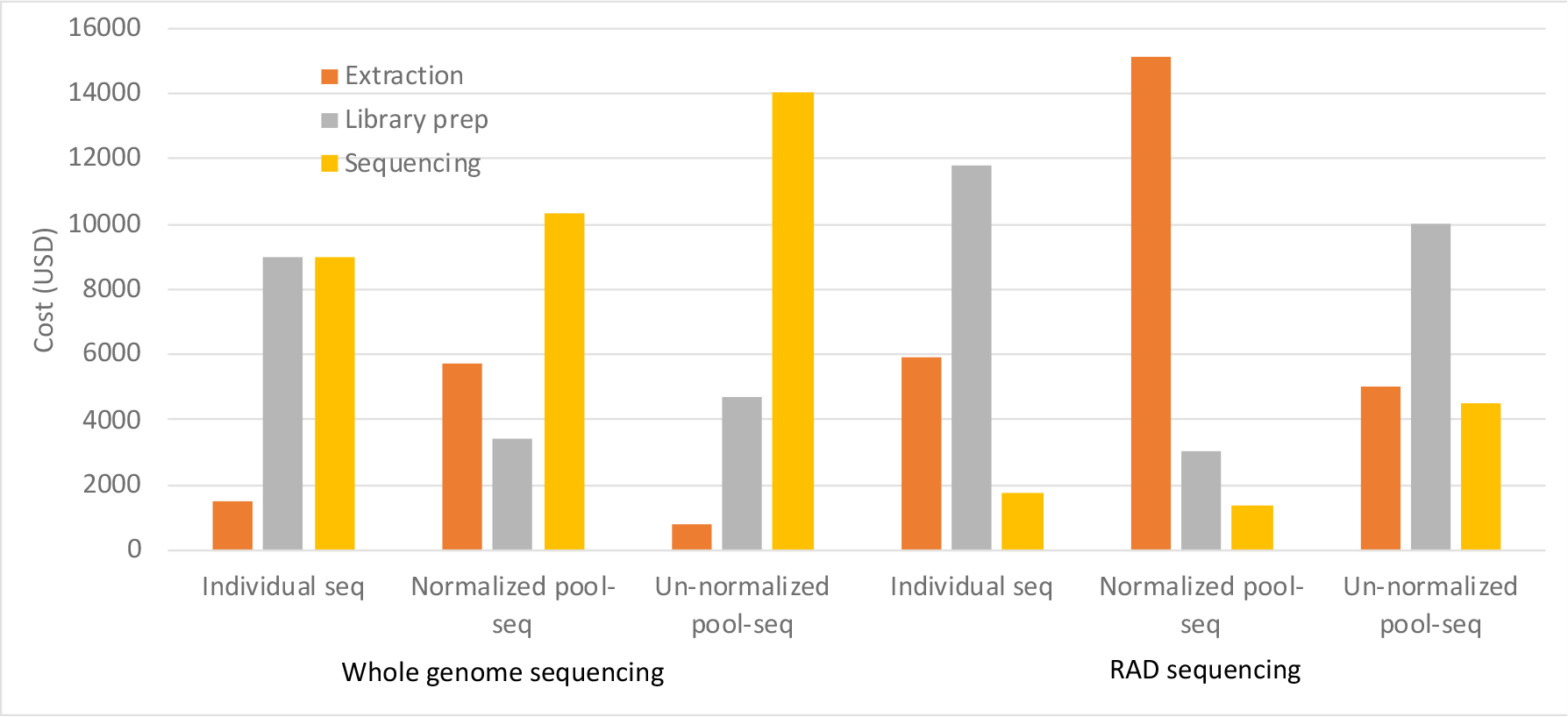
A distribution of costs for a given budget when using different DNA extraction, library construction and sequencing methods.

## Discussion

In this study, we report the first use of empirical measurements of experimental error in pooling DNA from multiple individuals for the purpose of evaluating the power of GWAS for detecting QTLs of colony-level traits. By evaluating the use of pool-seq vs. individual sequencing and WG-seq vs. RAD-seq, our study provides a broad overview on the experimental errors and biases in allele frequency estimations. Such pool-seq experimental biases can result from DNA quantification errors before normalization, pipetation errors, and various biases and artifacts in library construction and sequencing. We quantified the correspondence between allele frequencies in the individual genotypes and the pool-seq data by their correlation. Using simulations, we estimated, for the first time, the statistical power for detecting a QTL of a colony-level trait by taking into consideration the additive effects of environmental noise to that of the experimental errors.

WG-seq of pools provides considerably more accurate allele frequency estimates than RAD-seq of pools. The correlation between allele frequencies in individuals and in pool-seq was *rho*=0.89 for WG-seq and *rho*=0.63 for RAD-seq. Surprisingly, the correlation was not affected by either pool size (30 vs. 10 pooled individuals) or normalization following DNA extraction. The weaker correlation with the pools for RAD-seq relative to WG-seq may be due to RAD-seq artifacts. Such artifacts may be, for example, polymorphism at restriction sites resulting in alleles that are not cleaved and sequenced, which may bias allele frequency estimation for adjacent SNPs. Our study also revealed what appears to be a technical artifact in pool-seq. For low frequency alleles (<10% of the colony genotypes), pool-seq misses the rare allele completely. That is, no reads were sequenced from the rare allele in the pooled samples. This cannot be explained simply by low sequencing depth, since the pool had a sequencing depth of 30X on average and no less than 10X per locus. At 10X, the probability of not obtaining any reads for an allele with 10% allele frequency is only 0.9^9 = 0.39. Thus, we conclude that technical artifacts and biases in the pool-seq procedure lead to missed rare alleles in almost all such loci. It should also be noted that pooling methods are disadvantageous when genotyping dominant/recessive markers. In such cases, pool-seq masks heterozygosity whereas individual genotyping enables distinguishing between two heterozygotes and a recessive homozygote.

Evaluation of QTL detection power in GWAS revealed that as the environmental noise level rises, the success rate of identifying a colony-level QTL drops dramatically. Sequencing more individuals per colony can compensate for this reduced success rate, but only to a limited extent. When using 50 colonies, the power of pool-seq is equivalent to the power of a sequencing a small number of individual samples - between four and ten individuals per colony, depending on the environmental noise. However, the reduced costs of pool-seq allows for increasing the number of colonies in the study from 50 to 156 within the same budget when doing WG-seq, and to 1000 colonies when doing RAD-seq. Thus, pool-seq has two opposite effects: reducing the accuracy of allele frequency estimation for each colony, while increasing the number of colonies. The combined result is a dramatic increase in detection power when using pool-seq. For example, at a noise level of 1, sequencing six individuals per colony, for 50 colonies, gives a detection power of 58.4%, while pool-seq of 150 colonies increases the power to 92.3%.

These results demonstrate that using colony pools facilitate substantial power increase within a given budget. In general, it would be our recommendation to use pool-seq. In some scenarios, individual sequencing may be preferable. For example, if the number of sampled colonies is limited then individual sequencing would provide greater power (e.g. sequencing six or more individuals gives greater power than pool-seq if only 50 colonies are available, at noise level 1 in Fig. 2). Individual genome sequencing may also be desirable when there are additional data studied at the individual level, such as RNA sequencing for the purpose of expression QTL (eQTL) analysis (Majewski & Pastinen, 2011).

Apart from comparing between WG-seq and RAD-seq and between different pool sizes, we also evaluated in our study an alternative and an even more cost effective method of pooling. We show that extracting DNA from a pool of individuals gives similar accuracy as normalized pooling of individually extracted DNA. This is in spite of significant size variation in this species, with up to three-fold difference between the largest and smallest individuals. This may be because the experimental noise in pool-seq is on the same order of magnitude, or greater, than the size variation. The use of non-normalized pooling allows reducing DNA extraction costs as well as labor efforts.

Previous studies also investigated potential biases in pool-seq. Futschik and Schlötterer (2010) evaluated the effect of unequal representation of individual DNA in pools. However, this study was not based on empirical measurements of the biases in representation. Another study by Van Tassell et al. (2008) empirically evaluated the accuracy of pool-seq combined with RAD-seq. Van Tassell et al. (2008) assessed the concordance of the genotypes of 66 individually sequenced samples with allele frequencies estimated by sequencing their pooled DNA. They reported a correlation coefficient *rho* = 0.67, which is similar to our *rho* = 0.68-0.75. However, neither studies investigated QTL detection power in the context of GWAS, when using pool-seq compared to individual genotyping.

## Conclusions

We conclude that for a given budget, pool-seq is preferable over individual sequencing for maximizing the QTL detection power in GWAS of colony level phenotypes. However, selection of library construction and sequencing approaches should consider the expected environmental noise, availability of samples, and manpower. We also conclude that when using pool-seq, sequencing whole genomes is preferable over sequencing reduced representation libraries and that, at least in social insects’ colonies, increasing the size of the sequenced pool is of little advantage. We also found that sequencing un-normalized pools may be of great advantage as it increases the number of colonies that can be analyzed, the power of GWAS, and reduces labor efforts dramatically.

## Data Accessibility statement

All sequence data were deposited to the NCBI SRA data base, accession number: PRJNA494296.

## Acknowledgement

E.P. was supported by Israel Science Foundation Grants no. 646/15, 2140/15, and 2155/15 and US-Israel Binational Science Foundation Grant no. 2013408.

## References

Alexander, D. H., Novembre, J., & Lange, K. J. G. r. (2009). Fast model-based estimation of ancestry in unrelated individuals.

Andrews, K. R., Good, J. M., Miller, M. R., Luikart, G., & Hohenlohe, P. A. J. N. R. G. (2016). Harnessing the power of RADseq for ecological and evolutionary genomics. 17(2), 81.

Baird, N. A., Etter, P. D., Atwood, T. S., Currey, M. C., Shiver, A. L., Lewis, Z. A.,… Johnson, E. A. J. P. o. (2008). Rapid SNP discovery and genetic mapping using sequenced RAD markers. 3(10), e3376.

Bastide, H., Betancourt, A., Nolte, V., Tobler, R., Stobe, P., Futschik, A., & Schlotterer, C. (2013). A genome-wide, fine-scale map of natural pigmentation variation in Drosophila melanogaster. PLoS Genet, 9(6), e1003534. doi:10.1371/journal.pgen.1003534

Benjamini, Y., & Hochberg, Y. J. J. o. t. r. s. s. S. B. (1995). Controlling the false discovery rate: a practical and powerful approach to multiple testing. 289–300.

Catchen, J., Hohenlohe, P. A., Bassham, S., Amores, A., & Cresko, W. A. J. M. e. (2013). Stacks: an analysis tool set for population genomics. 22(11), 3124–3140.

Catchen, J. M., Amores, A., Hohenlohe, P., Cresko, W., & Postlethwait, J. H. J. G. G., genomes, genetics. (2011). Stacks: building and genotyping loci de novo from short-read sequences. 1(3), 171–182.

Catchen, J. M., Hohenlohe, P. A., Bernatchez, L., Funk, W. C., Andrews, K. R., & Allendorf, F. W. J. M. E. R. (2017). Unbroken: RADseq remains a powerful tool for understanding the genetics of adaptation in natural populations. 17(3), 362–365.

Darvasi, A., & Soller, M. (1994). Selective DNA pooling for determination of linkage between a molecular marker and a quantitative trait locus. Genetics, 138(4), 1365–1373.

Dunnington, E. A., Haberfeld, A., Stallard, L. C., Siegel, P. B., & Hillel, J. (1992). Deoxyribonucleic acid fingerprint bands linked to loci coding for quantitative traits in chickens. Poult Sci, 71(8), 1251–1258. doi:10.3382/ps.0711251

Eyer, P., Seltzer, R., Reiner-Brodetzki, T., & Hefetz, A. (2017). An integrative approach to untangling species delimitation in the Cataglyphis bicolor desert ant complex in Israel. 115, 128–139.

Futschik, A., & Schlötterer, C. J. G. (2010). The next generation of molecular markers from massively parallel sequencing of pooled DNA samples. 186(1), 207–218.

Gautier, M., Foucaud, J., Gharbi, K., Cézard, T., Galan, M., Loiseau, A.,… Estoup, A. J. M. E. (2013). Estimation of population allele frequencies from next-generation sequencing data: pool-versus individual-based genotyping. 22(14), 3766–3779.

Giovannoni, J. J., Wing, R. A., Ganal, M. W., & Tanksley, S. D. J. N. A. R. (1991). Isolation of molecular markers from specific chromosomal intervals using DNA pools from existing mapping populations. 19(23), 6553–6568.

Korol, A., Frenkel, Z., Cohen, L., Lipkin, E., & Soller, M. (2007). Fractioned DNA pooling: a new cost-effective strategy for fine mapping of quantitative trait loci. Genetics, 176(4), 2611–2623. doi:10.1534/genetics.106.070011

Leniaud, L., Heftez, A., Grumiau, L., & Aron, S. J. B. J. o. t. L. S. (2011). Multiple mating and supercoloniality in Cataglyphis desert ants. 104(4), 866–876.

Lowry, D. B., Hoban, S., Kelley, J. L., Lotterhos, K. E., Reed, L. K., Antolin, M. F., & Storfer, A. J. M. e. r. (2017a). Breaking RAD: An evaluation of the utility of restriction site-associated DNA sequencing for genome scans of adaptation. 17(2), 142–152.

Lowry, D. B., Hoban, S., Kelley, J. L., Lotterhos, K. E., Reed, L. K., Antolin, M. F., & Storfer, A. J. M. e. r. (2017b). Responsible RAD: striving for best practices in population genomic studies of adaptation. 17(3), 366–369.

Majewski, J., & Pastinen, T. J. T. i. G. (2011). The study of eQTL variations by RNA-seq: from SNPs to phenotypes. 27(2), 72–79.

McKenna, A., Hanna, M., Banks, E., Sivachenko, A., Cibulskis, K., Kernytsky, A.,… Daly, M. J. G. r. (2010). The Genome Analysis Toolkit: a MapReduce framework for analyzing next-generation DNA sequencing data.

McKinney, G. J., Larson, W. A., Seeb, L. W., & Seeb, J. E. J. M. e. r. (2017). RAD seq provides unprecedented insights into molecular ecology and evolutionary genetics: comment on Breaking RAD by Lowry et al.(2016). 17(3), 356–361.

Meaburn, E., Butcher, L. M., Liu, L., Fernandes, C., Hansen, V., Al-Chalabi, A.,… Schalkwyk, L. C. J. B. g. (2005). Genotyping DNA pools on microarrays: tackling the QTL problem of large samples and large numbers of SNPs. 6(1), 52.

Michelmore, R. W., Paran, I., & Kesseli, R. J. P. o. t. n. a. o. s. (1991). Identification of markers linked to disease-resistance genes by bulked segregant analysis: a rapid method to detect markers in specific genomic regions by using segregating populations. 88(21), 9828–9832.

Parchman, T. L., Gompert, Z., Mudge, J., Schilkey, F. D., Benkman, C. W., & Buerkle, C. A. J. M. e. (2012). Genome-wide association genetics of an adaptive trait in lodgepole pine. 21(12), 2991–3005.

Peterson, B. K., Weber, J. N., Kay, E. H., Fisher, H. S., & Hoekstra, H. E. (2012). Double digest RADseq: an inexpensive method for de novo SNP discovery and genotyping in model and non-model species. PLoS One, 7(5), e37135. doi:10.1371/journal.pone.0037135

Plotsky, Y., Cahaner, A., Haberfeld, A., Hillel, J., Lavi, U., & Lamont, S. J. A. G. (1993). DNA fingerprint bands applied to linkage analysis with quantitative trait loci in chickens. a. 24(2), 105–110.

Privman, E., Cohen, P., Cohanim, A. B., Riba-Grognuz, O., Shoemaker, D., & Keller, L. J. M. E. (2018). Positive selection on sociobiological traits in invasive fire ants.

Reiner-Brodetzki, T., Inbar, S., Cohen, P., Hefetz, A., & Privman, E. Social polymorphism in Cataglyphis niger. In preperation.

Schlötterer, C., Tobler, R., Kofler, R., & Nolte, V. J. N. R. G. (2014). Sequencing pools of individuals—mining genome-wide polymorphism data without big funding. 15(11), 749.

Van Tassell, C. P., Smith, T. P., Matukumalli, L. K., Taylor, J. F., Schnabel, R. D., Lawley, C. T.,… Sonstegard, T. S. J. N. m. (2008). SNP discovery and allele frequency estimation by deep sequencing of reduced representation libraries. 5(3), 247.

